# Dimethylarsenic acid (DMA) accumulation positively correlates with realgar-induced subchronic toxicity in rats

**DOI:** 10.1101/398206

**Authors:** Yan Yi, Shuangrong Gao, Jing Xia, Yong Zhao, Chunying Li, Yushi Zhang, Lianmei Wang, Chen Pan, Suyan Liu, Jiayin Han, Aihua Liang, Ji Shen

## Abstract

The toxicity of realgar depends largely on different arsenic species accumulation and distribution in the body. Here, after continuous oral administration of different doses of realgar for 90 days and subsequent 60-day withdrawal period, clinical observations, food consumption, body weights, blood biochemistry, hematology, and histomorphological examination of rats were performed. Realgar 40mg·kg^−1^·d^−1^ and 170 mg·kg^−1^·d^−1^ of realgar (which is equivalent to 40-fold and 100-fold the maximum clinical dose, respectively) can cause toxicity in rats, including degreased body weight, peripheral blood neutrality abnormal ratio of granulocytes and lymphocytes, hypercoagulability of the blood, liver and kidney tissue damage, liver and kidney may be the main toxic target organs of realgar. The no observed adverse effect level (NOAEL) dose is 10 mg·kg^−1^. At the same time, the content and distribution of arsenic species in tissues were determined. The content of total arsenic (tAs) and Dimethylarsenic acid (DMA) in the tissues of the realgar group was significantly higher than those of the control group. After 60 days of discontinuation, the DMA content in the realgar group decreased, but it was still higher than that in the control group, and liver and kidney damage occurred during the administration period basically returned to normal. Therefore, the authors speculated that when the DMA content in the tissue exceeds a certain range, liver and kidney toxicity will be induced. However, when the DMA content is lower than the above threshold after drug withdrawal, the liver and kidney lesions can return to normal.

## Introduction

Realgar is mineral medicine containing arsenic, the main component of which is As4S4 and its content is more than 90%. As a traditional Chinese medicine, realgar has a widely usage in clinic for thousands of years. Currently, more than 30 realgar preparations are still in use in China [1]. Realgar is effective in the treatment of various diseases such as infectious diseases (malaria, syphilis, parasitic infections), gastrointestinal diseases (ulcers), nervous system diseases (convulsions), rheumatic diseases and skin diseases (psoriasis) [2-4]. Modern researches also show that realgar has a significant anticancer effect in the clinic, especially on hematological cancer. Since the treatment of leukemia in the 1970s, more and more studies have shown that realgar has a potential therapeutic effect on various cancers. There is a growing understanding of the mechanism of action of realgar as an anti-tumor agent [5]. For example, realgar and realgar-containing formulations are effective and low toxicity in inducing apoptosis or differentiation of acute promyelocytic leukemia (APL) cells. However, realgar has been known as poisons since ancient times because arsenic is a well-known toxic substance. It was found that arsenic can mediate oxidative stress, induce hepatic apoptosis, cause chromosomal abnormality, alter growth factors expression, inhibit DNA repairing, increase the secretion of inflammatory factors and up regulate apoptosis related proteins, so that exposure to arsenic may induce liver disease, cardiovascular disease, nephrotoxicity, neurotoxicity and carcinogenicity [6-8].

The poor solubility of realgar is a barrier to its bioavailability, and long-term high-dose administration of realgar is necessary for clinical complete remission of cancer, but its potential toxicity limits its use. The long-term accumulation of toxic arsenic compounds in the body can cause abnormal tissue structure and function of the body [9-11]. Due to the known toxicity characteristics of arsenic [12-14], the long-term toxicity of realgar has always been a concern. However, it is unscientific to treat the toxicity of realgar and arsenic equally. Lu et al [15] compared the long-term renal toxicity of realgar, sodium arsenite (1/100 of the arsenic level) and sodium arsenate (1/50 of the arsenic level) in mouse. The results showed that the renal pathological lesions caused by sodium arsenite and sodium arsenite were more severe than that caused by realgar. Sodium arsenate and sodium arsenite induced the expression of heme oxygenase-related gene heme oxygenase-1, causing kidney damage, and realgar has no effects on this. Thus, it suggests that the toxicity of arsenic is closely related to its chemical form. There are studies that have demonstrated that if long-term ingestion of realgar, the total arsenic content in rat tissues will increase significantly and will cause damage to the liver and kidney of rats [16-17]. However, there are still few studies on its metabolic characteristics and exact toxic substances and toxic effects.

It is known that the metabolism of realgar is different from that of other substances containing arsenic, and the accumulation of As species may also be different. After administration of realgar, the arsenic metabolism is converted to various arsenic compounds. The toxicity of arsenic depends largely on its chemical structure, oxidation state and metabolic pathways. Various arsenic metabolites exhibit differing toxicities. For example, IC50 values of the main arsenic species is: AsIII < AsV< MMA≤DMA < AsC< AsB [18-20]. Thus, the quantity of each of these forms accumulated in tissues could be an important biomarker for evaluating arsenic toxicity. A large number of experiments have been conducted to study the changes in the total arsenic content of realgars entering the body, but there are few studies on their metabolism and toxicity profiles in the body. The association between the accumulation and distribution of arsenic species and the long-term toxicity of realgar is still not fully explored. The speciation information will contribute to improve our understanding of realgar metabolic transformation and toxicity mechanism. In this study, high performance liquid chromatography-inductively coupled plasma mass spectrometry (HPLC-ICP-MS) [21] was used to analyze arsenic species and the relationship between arsenic metabolism and realgar toxicity after long-term oral administration of realgar was evaluated. With the results we can figure out an underlying toxic substance basis and the main toxicity target organ of realgar, so that to provide scientific basis for further study the mechanism of the toxicity of realgar, clinically rational application of male-containing drugs, and their in-depth development.

## Materials and methods

### Reagents and Instruments

Realgar which derived from Shimen realgar mine (Hunan, China) was purchased from Shanghai Huayu Chinese Herbals Co., Ltd. (Shanghai, China) and processed in Shanghai Fengbang traditional Chinese Herbal Pieces factory (Shanghai, China). Blood cytology reagent was got from Yantai Zhuoyue biotechnology Co., Ltd (Shandong, China), and hematology were measured by Nihon Kohden (MEK-6318K) automatic blood cell analyzer (Tokyo, Japan). The coagulation test kits were bought from Beijing Shidi Science & Technology Instrument Co., Ltd (Beijing, China) and tested with C2000-4 Coagulation Aanlyzer produced by Beijing Precil Instrument Co., Ltd (Beijing, China). Urine routine testing were analyzed by urinary test papers in Bayer 50 Urine Analyzer (Leverkusen, Germany). Blood biochemical analysis kits were obtained from BioSino Biotechnology & Science Inc. (Beijing, China) and the relevant tests were carried out on Dimension AR biochemistry analyzer (Deerfield, Illinois, USA). Medica easylyte plus Na/K/Cl analyzer (Bedford, USA) was used to measure electrolyte. Electrolytes analysis reagent were purchased from Electra-Med Corporation (Michigan, USA). Reagents such as CMC-Na, Formaldehyde, ethanol, methanol, acetonitrile and hydrochloric acid were purchased from Sigma-Aldrich (St. Louis, Missouri, USA). All pathology instruments were from Leica biosystems (Vienna, Vienna, Austria) except the microscope which comes from Olympus Imagine (Tokyo, Japan). CEM Mars5 microwave (Matthews, North Carolina, USA) instrument was used for Arsenic extraction and wet digestion. The HPLC-ICP-MS system consisted of a Perkin Elmer OPTIMA-DV5300 (Waltham, Mass, USA) and an Agilent 7700ce ICP-MS systems (California, USA).

### Ethical statement

The entire animal experiment was conducted the recommendations in the Guide for the Care and Use of Laboratory Animals of the Institute of Chinese Materia Medica, China Academy of Chinese Medical Sciences. The protocol was approved by the Committee on the Ethics of Animal Experiments of the Institute of Chinese Materia Medica, China Academy of Chinese Medical Sciences (Permit Number:20102016).

### Animals

80 male and 80 female Wistar rats, approximately seven-week-old, were obtained from Vital River, a Charles River Company, Beijing, China. All rats were fed according to animal management standards of People’s Republic of China (temperature: 20-24°C, humidity:45-70%, ventilation:15 times per hour and 12h/12h light/dark cycle). Rats recovered in the laboratory for 1 week on free access to fixed-formula rat granule feed and pure water prior to any procedures. According to the ICH M3 sub chronic toxicity test guidelines [22], we selected an animal dosing cycle of 90 days and a recovery period of 60 days. When the animal weighed ≤ 350g, 5 rats per cage were fed. When the animal weighed more than 350g and less than 600g, 3 rats per cage were fed. When the animal weighed ≥ 600g, 2 rats per cage were fed. We checked the animal health condition every day. There were no unexpected deaths during our experiments. We first anesthetized the animals and then sacrificed the animals by exsanguinating abdominal aortic at the endpoint of the study. In addition, we used humane endpoints for animals when they were showing signs of moribund conditions including hypothermia and 20% weight loss. No mice reached this moribund state during the experiment. We carried out a single-cage feeding and provided supplemental heat and massaged when the animals have debilitating symptoms.

### Toxicity test of rats orally realgar for 90 days

According to Chinese Pharmacopoeia 2015 Edition, the maximum clinical dosage of realgar is 1.7 mg·kg^−1^/d. In this study, the four doses set were 0 mg·kg^−1^·d^−1^, 10 mg·kg^−1^·d^−1^, 40 mg·kg^−1^·d^−1^and 170 mg·kg^−1^·d^−1^ were set, which is equivalent 6-fold, 24-fold and 100-fold of 1.7 mg·kg^−1^, respectively. Thus, the rats were randomly delivered as 4 groups, 18 male rats and 18 female rats in each group. Rats in the 0 mg·kg^−1^·d^−1^ group were orally administrated with 0.3% Carboxymethyl Cellulose Sodium (CMC-Na) solution each day, and realgar was dissolved with 0.3% CMC-Na ultrapure water. Rats administration volume is 5 ml/kg, and the other three realgar groups’ rats were given 2 mg·ml^−1^, 8 mg·ml^−1^, 34 mg·ml^−1^realgar-0.3% CMC-Na solution respectively. After 60 and 90 days of administration, each group of five male rats and female rats were sacrificed. Finally, each of the remaining groups of rats ceased drug treatment and were fed for 60 days and then were sacrificed on the 30th and 60th days of discontinuation, respectively. During the experiment, the general condition of the rats was recorded daily, including gait, general activity, hair, stool, urine, and so on. Body weight of all rats and food consumption per cage were measured weekly.

Before each sacrifice, rats were individually placed into metabolic cages to collect 24 h-urine. Urine was qualitatively analyzed and indicators included urine glucose (GLU), ketone body (KET), bilirubin (BIL), pH, specific gravity (SG), occult blood (BLO), protein (PRO), urobilinogen (URO), leukocytes (Leu). Thereafter, rats were anesthetized with phenobarbital sodium 30 mg·kg^−1^ i.p, then the blood of each rat was taken from abdominal aorta to detect the hematology (including Red blood cell count (RBC), white blood cell count (WBC), hemoglobin (HGB), Red blood cell specific volume (HCT), blood platelet number (PLT), mean corpuscular volume (MCV), mean corpuscular hemoglobin (MCH), mean corpuscular hemoglobin concentration (MCHC), leukocyte differential count and reticulocyte), coagulation function (prothrombin time (PT) and activated partial thromboplastin time (APTT)), biochemistry(serum total protein(TP), albumin(ALB), aspartate transaminase(AST), alanine transaminase(ALT), alkaline phosphatase(ALP), γ-glutamyl-transferase(γ-GT), creatine kinase(CK), blood urea nitrogen(BUN), creatinine(CRE), glucose(GLU), total bilirubin(TBIL), total cholesterol(T-CHO), triglyceride(TG), and electrolytes (Na^+^, K^+^ and Cl^−^).

Next, autopsy was performed. Rats’ all organs including brain, pituitary, salivary gland, thymus, lung with bronchi, heart, liver, spleen, kidney, stomach, adrenal gland, testicle, prostate, seminal vesicle, ovary, uterus was all removed together and weighted. Relative organ weight were calculated as follows: Relative Organ Weight(%) (ROW)=[organ weight(g)/body weight before death(g)]×100%. After weighting, all organs were fixed with 10% formalin, embedded in paraffin, sectioned, stained with HE (Haematoxylin & Eosin) stained, and then observed under a microscope for histomorphology. Histology of the liver, kidney and other major organs was scored from mild to severe according to pathological changes. The degree of liver tissue pathological changes was scored using 3 grades [23]: Grade 0–no obvious lesions; Grade1– the structure of the hepatic lobule is still normal, with turbid swelling, eosinophilic degeneration or steatosis, and scattered necrosis; Grade 2– Lobular structure of the liver is unclear, showing focal necrosis with inflammatory cell infiltration. The degree of renal tubular injury was scored using 3 grades [24]: Grade 0–kidney structure is normal; Grade 1 – the brush border of tubular epithelial cells was lost, cells were slightly swollen, vacuolar degeneration, a small amount of protein tube in the lumen, mild interstitial edema, and a small amount of inflammatory cell infiltration; Grade 2 – renal tubular epithelial cells are swollen, with loose cytoplasm, vacuolar degeneration, and even necrosis, loss, visible cells tube type and protein tube type, obvious interstitial edema.

### Rat biological sample collection and pretreatment

At each sacrifice, rats’ blood, urine, feces, liver, kidney and brain of rats were obtained to test the arsenic species therein. Determination of arsenic species in blood is usually done with 2 ml of 10% EDTA-K2 anticoagulant. 5 ml of urine and all feces in the metabolic cage were required to detect the exact arsenic levels in urine and feces. At least 1g of organs (liver, kidney and brain) should be kept to measure the level of arsenic in it. All samples were kept at −80°C before the detection of arsenic species. Before testing, the liver, kidney and brain were homogenized and the excrement was dried and crushed. We respectively weighed the above samples, urine and plasma 0.5g each, then put them into the microwave digestion tank, add 4mL nitric acid and 1mL hydrochloric acid for digestion. Then the samples were taken out for cooling, transferred to a 50mL polytetrafluoroethylene volumetric flask while the digestion tank is washed several times with deionized water and the wash liquid were poured into the measuring bottle. After that, we added deionized water until the liquid level reached 50ml mark and mix the liquid evenly to get the final samples.

### Analysis of arsenic species accumulation and distribution in biological samples by HPLC-ICP-MS

Various forms of arsenic standard stock solutions are prepared with ultrapure water and diluted to the working liquid on the day of the test. The samples are acidificated with hydrochloric acid (HCL) and then added to an ion exchange resin chromatography column, which are then eluted with to give an eluate. Since only arsenic species such as MMA and DMA can adsorb to the column, AsIII and AsV can be separated from total arsenic through the column. The column is then separated by adding ultrapure water and MMA and DMA are obtained by collecting the dissociation liquid. Next, dilute HCL eluate containg AsIII and AsV is passed through an anion strong basic exchange column and then subsequently AsV is absorbed into the column, thus obtaining AsIII by collecting the eluate. In order to get AsV, we add HCl to the anion exchange column to dissolve AsV to obtained AsV HCl solution. Finally, all AsIII AsV MMA and DMA samples were detected by ICP-MS except that the fecal samples were determinate by inductively coupled plasma atomic emission spectrometry (ICP-OES). Instrumental conditions for ICP-MS were as follows: RF power,1500W; Sampling depth,8.0mm; Carrier gas flow rate, 0.9L·min^−1^, Carrier gas compensation air flow rate,0.3L·min^−1^; the analysis model, quantitative analysis (quantitative analysis model), Integration time, 2s; repeat 3 times. ICP-OES detection mas set as: power,1300W; Atomizing gas flow, 0.8L ·min^−1^); Auxiliary gas flow, 0.2L·min^−1^, Plasma gas flow, 15L·min^−1^, Peristaltic pump flow rate, 15mL·min^−1^; Integration time,1∼20s; Radial observation, line wavelength 188.979nm.

### Statistical analysis

If not indicated otherwise, data are expressed as arithmetic means(M) ± standard error of mean (S.E.M). The data of body weights, food consumption, organ weight, urology, hematology and blood biochemistry values of male and female rats were analyzed separately. The arsenic content data of male and female rats were combined and analyzed. These data were performed using parametric one-way analysis of variance (ANOVA) followed by LSD (equal variances assumed) or Tamhane’s T2 (equal variances not assumed) multiple comparison test. The difference between each realgar-treated group and the control group were compared. Statistical differences in quantitative data were obtained by analysis of variance (ANOVA), qualitative urinary data and histopathological changes were statistically analyzed with Kruskal-Wallis H-test and Mann-Whitney U-test. Statistical analysis using SPSS 17.0 software. A *p*-value of <0.05 was considered to be statistically significant.

## Results and discussion

### Oral realgar caused a degrease in body weight in rats

After 90 days of administration, except the rats in 170 mg·kg^−1^·d^−1^ realgar group had depilated, while other animals were generally in good condition. In addition, from the 60th day to the 90th day, compared with the 0 mg·kg^−1^·d^−1^ control group, the body weight of female rats in the 40 mg·kg^−1^·d^−1^ realgar group was significantly decreased (p < 0.05). After discontinuation of treatment, the body weight of male rats in the realgar group showed a downward trend compared with the control, but there was no statistical difference (Fig 1). There was no difference in the average food consumption between the four groups during treatment and after discontinuation (data not shown). Since the weight-reduced rats returned to normal after drug withdrawal, we believe that this adverse reaction may be mainly due to the larger volume and the higher concentration of the drug, which affects the rats’ feeding and digestive functions, rather than the toxic effects of realgar.

**Fig 1.**
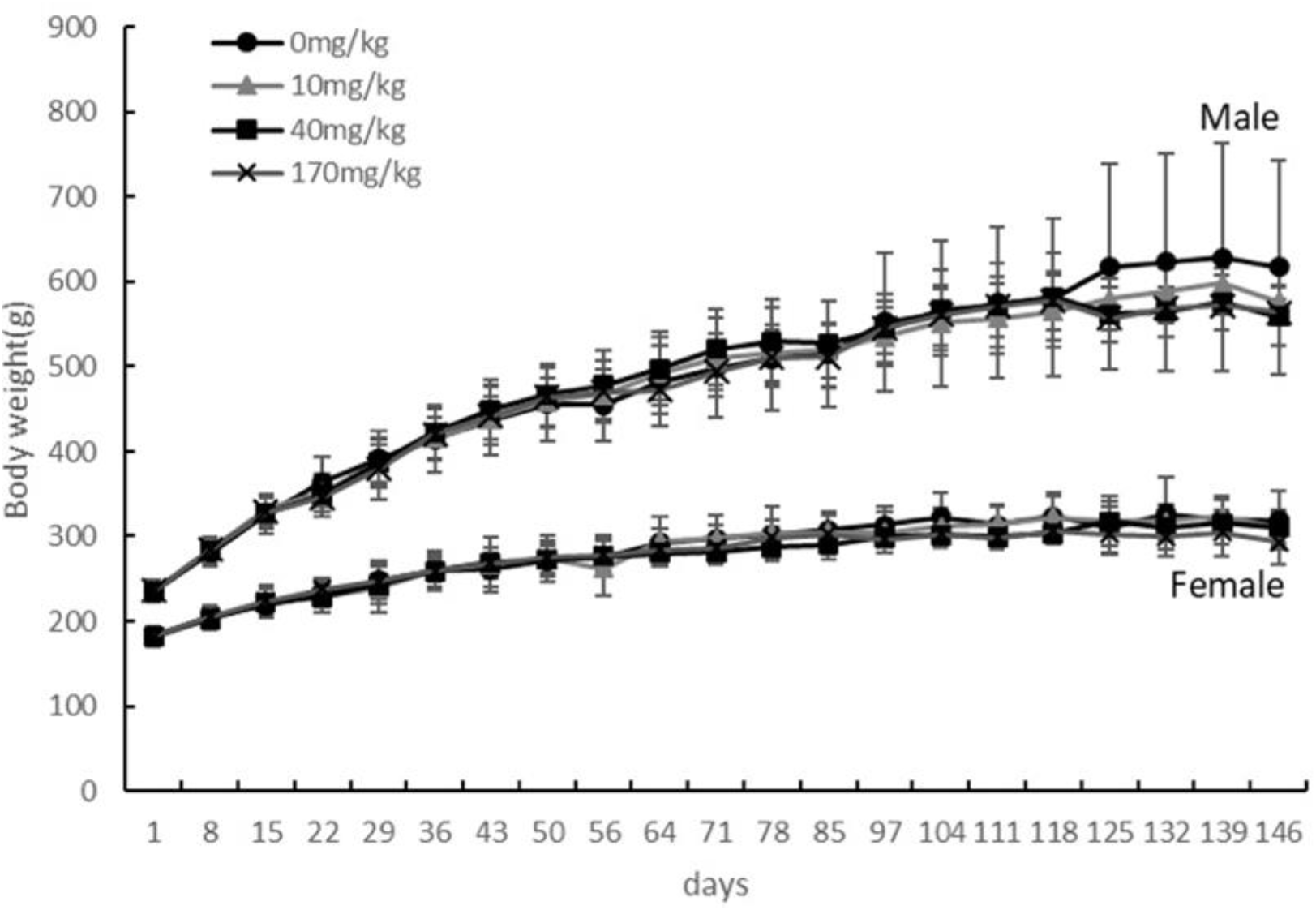
The effect of the oral realgar treatment on mean body weight in male and female rats throughout the whole experiment. Values are presented as mean ± S.E.M.

### Realgar may result in abnormal of peripheral blood neutrophils and lymphocytes ratios

After treatment with realgar for 90 days, compared with the control rats, in male and female rats of the realgar medium-dose group and high-dose group, lymphocyte ratio (Ly%) showed an upward trend, and neutrophilic granulocyte ratio (Gr%) showed a decreasing trend (*p*<0.05, *p*<0.01) (Table 1). We retrospectively reviewed the results of the 60-day study, with no apparent abnormalities in Ly% and Gr% (data omitted). After discontinuation of treatment, the values of Ly% and Gr% in the high-dose and medium-dose group of the realgar gradually returned to normal, and there was no difference from the control group 60 days after drug withdrawal (Table 2). Other than that, there were no obvious abnormal changes in other hematological parameters during realgar administration, and after 60 days of discontinuation of the drug, there were no delayed toxicities in the realgar hematological parameters. Therefore, long-term (90-day) use of large doses of realgar (100 times the maximum clinical dose) may have an effect on the immune function of rats. However, the specific impact on immune function needs to be further studied. After the realgar is no longer used, peripheral blood neutrophils and lymphocytes ratios of the rats returned to normal.

**Table 1.**
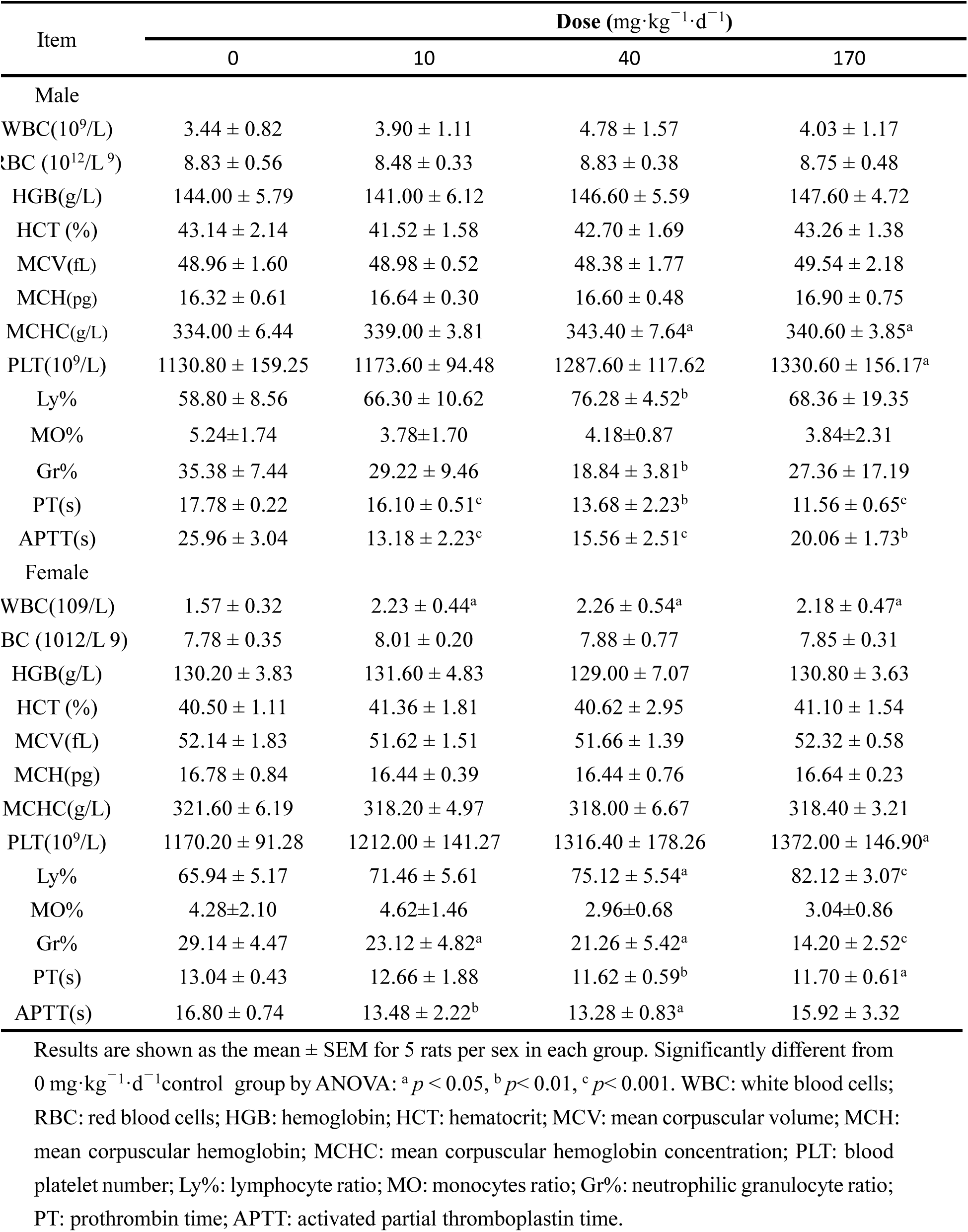
Hematological data for male and female rats orally administered realgar for 90 days(n=5/sex/group, 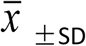)

**Table 2.**
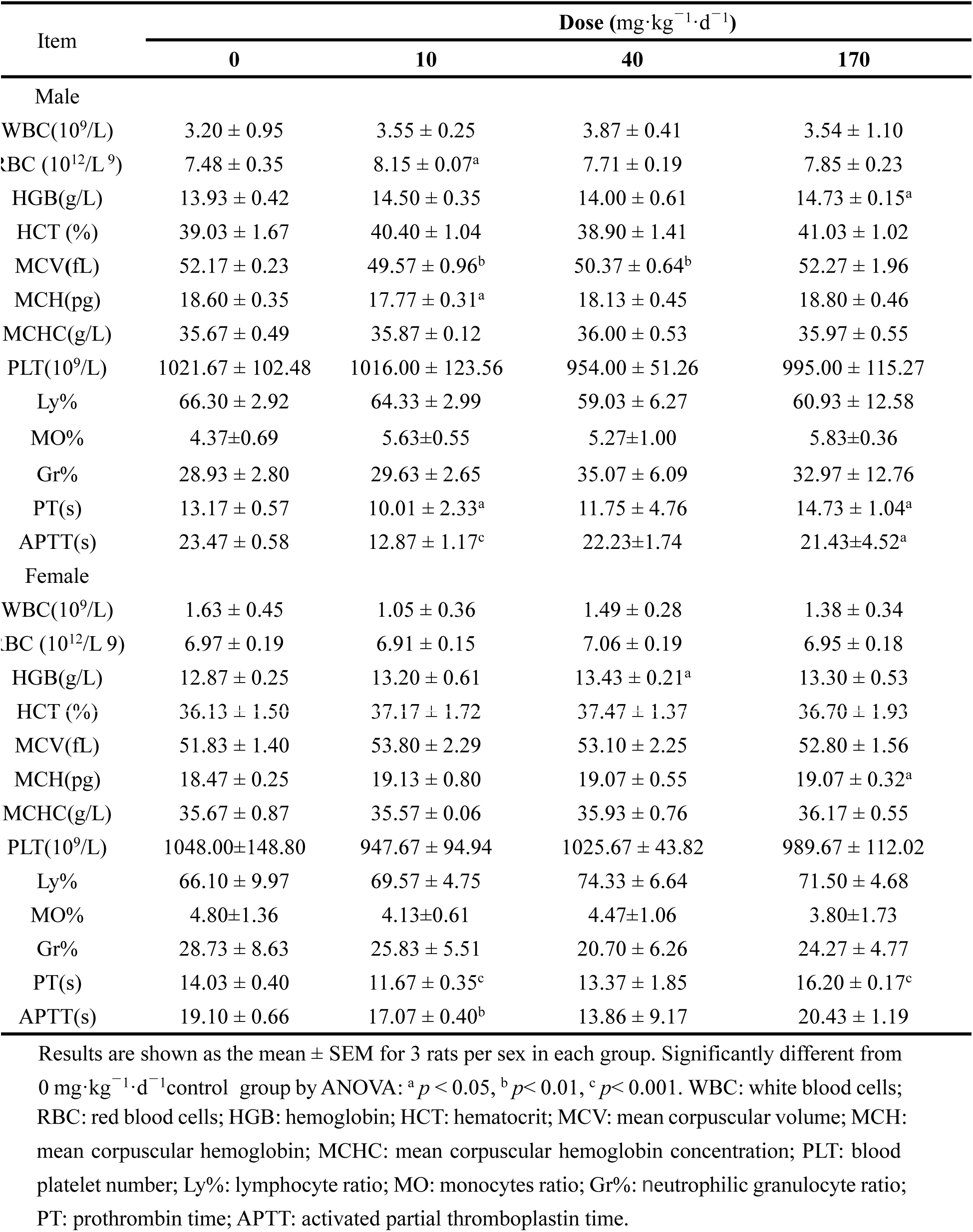
Hematological data for male and female rats after 60 days’ withdrawal(n=3/sex/group, 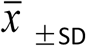)

### Realgar may cause hypercoagulability in rats

During the treatment period, compared with the rats in 0 mg·kg^−1^·d^−1^ control group, PT and APTT of animals (♂, ♀) in low, middle and high dose realgar groups significantly decreased (*p* <0.01), so long-term use of realgar can cause blood hypercoagulation. The hypercoagulable state of blood has the risk of thrombosis, which in turn causes damage to various systems of the body, in particular blood vessels and the heart. 60 days after drug withdrawal, the APTT and PT values (♂, ♀) of the realgar group gradually returned to normal (Table 1-2). It is reported that realgar can decrease the expression of apoptosis-regulating gene Bc1-2, down-regulate the expression of apoptosis protein (survivin), increase the expression of pro-apoptotic gene Bax protein, and promote the apoptosis of tumor cells, so as to achieve the purpose of treating acute promyelocytic leukemia (APL) [25]. However, thrombosis and bleeding complications are the main causes of early death of APL. So far, there is no report on whether realgar can cause hypercoagulability that may have side effects on the treatment of leukemia. Since our results are mouse-based and there are differences between humans and mice, we can only initially suspect that realgar will cause clinical hypercoagulability. Since the DMA in the blood of the realgar rats was significantly higher than that of the control group, whether or not the hypercoagulability state is related to the high level of DMA and its specific mechanism,all these requires further study.

### Realgar may damage kidney function and may also cause hypercholesterolemia and abnormal electrolyte levels

After 60 days of administration, urinary protein (PRO) and white blood cell count (WBC) in realgar groups were significantly higher than those in the control group (*p*<0.05). However, after 90 days of dosing, this condition did not worsen, and they gradually returned to normal after discontinuation. There are no other significant differences in urinalysis results (data not shown). 90 days after the administration of realgar, the blood urea nitrogen (BUN) values of rats in high-dose group (♂) and low-dose group (♀) increased significantly (*p* < 0.05) (Table 3). While after the 60 days of drug withdrawal, there was no obvious abnormality in BUN value in the realgar group. In addition, cholesterol (CHO) levels were significantly increased in the realgar group (♂) 60 days after administration (p < 0.05), and all the realgar group rats CHO until 90 days after dosing and 30 days after discontinuation values (♂, ♀) are still higher than control rats (*p* < 0.05). Serum BUN and urinary protein were significantly increased in rats, suggesting that renal function may be impaired. Compared with 0 mg·kg^−1^·d^−1^ control group, there was no significant difference in serum biochemical indexes in realgar groups’ rats after 60 days of withdrawal (Table 4). The result of blood electrolyte analysis showed that after 60 days of realgar administration, the K+ value of some realgar groups (♂, ♀) decreased (*p* < 0.05). Furthermore, after 90 days’ administration and 60 days after drug withdrawal, the K+ values of male rats in some realgar groups were also significantly lower than that of the control rats (*p* < 0.05). However, after 90 days of realgar treatment, the K+ value of female rats in high-dose realgar group was significantly higher than that of the control group (*p* < 0.01) (Table3-4). In short, long-term high-dose use of realgar has certain effects on rat cholesterol metabolism and blood electrolyte (K^+^) levels, but its specific causes and the relationship with arsenic levels in the body is still unclear.

**Table 3.** Serum biochemistry data for male and female rats orally administered realgar for 90 days (n=5/sex/group, 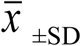)

| Item | Dose (mg·kg<sup>-1</sup>·d<sup>-1</sup>) |  |  |  |
| --- | --- | --- | --- | --- |
|  | 0 | 10 | 40 | 170 |
| **Male** |  |  |  |  |
| ALB(g/L) | 33.16 ± 1.96 | 34.26 ± 1.91 | 35.32 ± 2.83 | 35.16 ± 1.09<sup>a</sup> |
| ALP(IU) | 68.47 ± 14.99 | 69.61 ± 12.68 | 58.12 ± 15.43 | 52.88 ± 12.50 |
| ALT (U/L) | 44.80 ± 12.93 | 45.80 ± 17.38 | 35.60 ± 16.49 | 28.40 ± 3.91<sup>a</sup> |
| AST(U/L) | 112.60 ± 17.14 | 102.20 ± 65.86 | 81.60 ± 27.95<sup>a</sup> | 84.60 ± 18.93<sup>a</sup> |
| CHO(mmol/L) | 1.79 ± 0.15 | 2.23 ± 0.24<sup>b</sup> | 2.14 ± 0.21<sup>b</sup> | 1.89 ± 0.18 |
| CK(U/L) | 576.40 ± 185.31 | 101.00 ± 66.75<sup>c</sup> | 210.20 ± 266.90<sup>a</sup> | 341.20 ± 218.11 |
| CRE(umol/L) | 53.80 ± 4.02 | 49.20 ± 3.35<sup>a</sup> | 48.80 ± 2.17<sup>a</sup> | 57.20 ± 5.67 |
| γ-GT (U/L) | 0.45 ± 0.07 | 0.44 ± 0.20 | 0.55 ± 0.53 | 0.32 ± 0.18 |
| GLU(mmol/L) | 7.14 ± 0.83 | 7.68 ± 0.24 | 7.21 ± 1.34 | 7.48 ± 0.95 |
| BUN(mmol/L) | 7.38 ± 0.89 | 6.95 ± 1.31 | 7.06 ± 0.79 | 9.91 ± 2.21<sup>a</sup> |
| TBIL(umol/L) | 0.58 ± 0.56 | 1.46 ± 0.41<sup>a</sup> | 1.12 ± 1.14 | 1.44 ± 0.84<sup>a</sup> |
| TG(mmol/L) | 0.51 ± 0.23 | 0.66 ± 0.21 | 0.88 ± 0.31<sup>a</sup> | 0.99 ± 0.23<sup>b</sup> |
| TP(g/L) | 54.96 ± 3.26 | 56.92 ± 3.27 | 57.98 ± 4.20 | 57.26 ± 0.93 |
| Na(mmol/L) | 138.72 ± 0.78 | 139.78 ± 0.65<sup>a</sup> | 139.10 ± 1.45 | 138.72 ± 0.95 |
| K(mmol/L) | 4.25 ± 0.25 | 3.57 ± 0.30<sup>b</sup> | 3.87 ± 0.20<sup>a</sup> | 3.99 ± 0.39 |
| Cl(mmol/L) | 105.44 ± 1.79 | 105.96 ± 0.98 | 104.90 ± 1.39 | 104.04 ± 2.01 |
| **Female** |  |  |  |  |
| ALB(g/L) | 35.76 ± 2.36 | 33.04 ± 2.02<sup>a</sup> | 36.18 ± 1.88 | 34.50 ± 1.29 |
| ALP(IU) | 39.43 ± 6.77 | 43.19 ± 9.83 | 30.71 ± 6.16<sup>a</sup> | 33.09 ± 4.46 |
| ALT (U/L) | 35.40 ± 3.58 | 36.20 ± 4.21 | 38.00 ± 17.58 | 29.80 ± 7.01 |
| AST(U/L) | 84.40 ± 12.86 | 88.00 ± 17.45 | 89.40 ± 17.67 | 88.20 ± 7.22 |
| **CHO(mmol/L)** | 1.63 ± 0.26 | 1.35 ± 0.40 | 1.76 ± 0.23 | 1.92 ± 0.19<sup>a</sup> |
| **CK(U/L)** | 266.00 ± 144.56 | 114.20 ± 69.59<sup>a</sup> | 175.00 ± 123.36 | 348.80 ± 105.23 |
| **CRE(umol/L)** | 57.00 ± 4.85 | 66.20 ± 10.80 | 58.00 ± 8.72 | 56.00 ± 3.67 |
| **γ-GT (U/L)** | 0.44 ± 0.15 | 0.64 ± 0.20 | 0.45 ± 0.17 | 0.44 ± 0.10 |
| **GLU(mmol/L)** | 7.48 ± 0.70 | 7.36 ± 0.81 | 6.78 ± 0.41<sup>a</sup> | 6.28 ± 0.22<sup>b</sup> |
| **BUN(mmol/L)** | 8.34 ± 0.75 | 10.60 ± 2.24<sup>a</sup> | 9.19 ± 1.42 | 9.38 ± 1.33 |
| **TBIL(umol/L)** | -0.04 ± 0.71 | -0.48 ± 0.31 | -0.16 ± 0.46 | 0.18 ± 0.16 |
| **TG(mmol/L)** | 0.29 ± 0.06 | 0.21 ± 0.08 | 0.30 ± 0.11 | 0.36 ± 0.06 |
| **TP(g/L)** | 57.26 ± 3.43 | 52.44 ± 3.59<sup>a</sup> | 57.62 ± 3.98 | 55.82 ± 2.18 |
| **Na(mmol/L)** | 139.98 ± 0.43 | 140.10 ± 1.20 | 140.50 ± 0.84 | 140.98 ± 0.90<sup>a</sup> |
| **K(mmol/L)** | 3.13 ± 0.19 | 3.22 ± 0.20 | 3.09 ± 0.22 | 3.46 ± 0.15<sup>b</sup> |
| **Cl<sup>-</sup>(mmol/L)** | 105.96 ± 1.68 | 106.48 ± 1.17 | 106.56 ± 1.96 | 107.64 ± 1.41 |

Results are shown as the mean ± SEM for 5 rats per sex in each group. Significantly different from 0 mg·kg<sup>-1</sup>·d<sup>-1</sup> control group by ANOVA: <sup>a</sup> $p < 0.05$ , <sup>b</sup> $p < 0.01$ , <sup>c</sup> $p < 0.001$ . ALB: total albumin; ALP: alkaline phosphatase; ALT: alanine aminotransferase; AST: aspartate aminotransferase; CHO: cholesterol; CK: creatine kinase; CRE: creatinine; GGT: γ-glutamyl transpeptidase; GLU: glucose; TBIL: total bilirubin; TG: triglycerides; TP: total protein; BUN: blood urea nitrogen; Na<sup>+</sup>: sodium; K<sup>+</sup>: potassium; Cl<sup>-</sup>: chloride.

**Table 4.** Serum biochemistry data for male and female rats after 60 days’ withdrawal(n=3/sex/group, 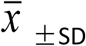

| Item | Dose (mg·kg<sup>-1</sup>·d<sup>-1</sup>) |  |  |  |
| --- | --- | --- | --- | --- |
|  | 0 | 10 | 40 | 170 |
| **Males** |  |  |  |  |
| **ALB(g/L)** | 32.37 ± 0.21 | 31.27 ± 0.65<sup>a</sup> | 31.43 ± 0.21<sup>b</sup> | 31.17 ± 0.64<sup>a</sup> |
| **ALP(IU)** | 48.93 ± 4.45 | 43.29 ± 6.92 | 56.14 ± 5.35 | 57.82 ± 6.73 |
| **ALT (U/L)** | 32.33 ± 0.58 | 33.00 ± 1.00 | 116.33 ± 107.83 | 36.33 ± 4.93 |
| **AST(U/L)** | 108.00 ± 20.66 | 81.67 ± 10.07 | 208.67 ± 155.76 | 105.67 ± 11.93 |
| **CHO(mmol/L)** | 2.09 ± 0.32 | 2.13 ± 0.07 | 1.61 ± 0.36 | 1.65 ± 0.16<sup>a</sup> |
| **CK(U/L)** | 509.67 ± 200.17 | 221.67 ± 106.28<sup>a</sup> | 242.00 ± 137.20 | 292.00 ± 147.25 |
| **CRE(umol/L)** | 51.67 ± 2.31 | 50.33 ± 3.79 | 50.67 ± 2.31 | 54.33 ± 7.09 |
| **γ-GT (U/L)** | 0.51 ± 0.11 | 0.50 ± 0.03 | 0.45 ± 0.11 | 0.58 ± 0.04 |
| **GLU(mmol/L)** | 6.92 ± 0.01 | 8.03 ± 1.22 | 6.92 ± 0.30 | 8.16 ± 1.27 |
| **BUN(mmol/L)** | 6.41 ± 0.67 | 5.12 ± 0.37<sup>a</sup> | 6.08 ± 0.69 | 6.00 ± 0.61 |
| **TBIL(umol/L)** | 5.70 ± 1.15 | 5.27 ± 0.95 | 5.47 ± 0.46 | 4.73 ± 0.51 |
| **TG(mmol/L)** | 0.57 ± 0.16 | 0.55 ± 0.18 | 0.50 ± 0.21 | 0.39 ± 0.14 |
| **TP(g/L)** | 54.13 ± 2.11 | 51.67 ± 1.14 | 51.90 ± 0.53 | 52.07 ± 1.93 |
| **Na(mmol/L)** | 139.70 ± 1.59 | 137.63 ± 5.38 | 141.13 ± 0.57 | 140.40 ± 1.01 |
| **K(mmol/L)** | 4.18 ± 0.20 | 4.25 ± 0.14 | 3.80 ± 0.15a | 4.01 ± 0.12 |
| **Cl(mmol/L)** | 107.87 ± 1.10 | 103.47 ± 4.60 | 106.97 ± 1.75 | 106.87 ± 1.18 |
| **Female** |  |  |  |  |
| **ALB(g/L)** | 33.00 ± 0.79 | 32.70 ± 1.91 | 33.83 ± 4.18 | 33.23 ± 1.80 |
| **ALP(IU)** | 40.84 ± 19.26 | 25.19 ± 4.90 | 28.81 ± 3.58 | 32.34 ± 6.48 |
| **ALT (U/L)** | 40.67 ± 5.69 | 34.00 ± 7.21 | 41.00 ± 4.58 | 40.00 ± 13.00 |
| **AST(U/L)** | 99.67 ± 14.57 | 86.33 ± 8.50 | 100.33 ± 32.33 | 105.00 ± 20.66 |
| **CHO(mmol/L)** | 1.32 ± 0.08 | 1.86 ± 0.46 | 1.63 ± 0.60 | 1.83 ± 0.44 |
| **CK(U/L)** | 288.00 ± 70.19 | 256.00 ± 38.16 | 298.33 ± 223.55 | 244.67 ± 111.85 |
| **CRE(umol/L)** | 56.67 ± 7.64 | 55.67 ± 4.04 | 53.00 ± 1.00 | 53.67 ± 1.53 |
| **γ-GT (U/L)** | 0.51 ± 0.14 | 0.52 ± 0.24 | 0.70 ± 0.26 | 0.91 ± 0.97 |
| **GLU(mmol/L)** | 7.23 ± 0.43 | 7.24 ± 0.74 | 7.23 ± 0.17 | 6.79 ± 0.41 |
| **BUN(mmol/L)** | 6.10 ± 1.07 | 6.44 ± 0.21 | 6.90 ± 0.22 | 6.74 ± 0.91 |
| **TBIL(umol/L)** | 4.43 ± 0.60 | 5.33 ± 1.37 | 5.73 ± 0.06<sup>a</sup> | 4.90 ± 0.82 |
| **TG(mmol/L)** | 0.34 ± 0.04 | 0.45 ± 0.19 | 0.44 ± 0.19 | 0.33 ± 0.05 |
| **TP(g/L)** | 53.53 ± 0.31 | 53.03 ± 1.53 | 54.10 ± 4.73 | 54.50 ± 1.47 |
| **Na(mmol/L)** | 140.80 ± 1.13 | 141.17 ± 1.08 | 140.87 ± 1.79 | 142.37 ± 0.15 |
| **K(mmol/L)** | 3.40 ± 0.17 | 3.21 ± 0.28 | 3.32 ± 0.15 | 3.19 ± 0.08 |
| **Cl(mmol/L)** | 107.73 ± 1.12 | 108.20 ± 1.35 | 107.17 ± 2.70 | 110.03 ± 0.74<sup>a</sup> |

Results are shown as the mean ± SEM for 3 rats per sex in each group. Significantly different from 0 mg·kg<sup>-1</sup>·d<sup>-1</sup> control group by ANOVA: <sup>a</sup> *p* < 0.05, <sup>b</sup> *p* < 0.01, <sup>c</sup> *p* < 0.001. ALB: total albumin; ALP: alkaline phosphatase; ALT: alanine aminotransferase; AST: aspartate aminotransferase; CHO: cholesterol; CK: creatine kinase; CRE: creatinine; GGT: γ-glutamyl transpeptidase; GLU: glucose; TBIL: total bilirubin; TG: triglycerides; TP: total protein; BUN: blood urea nitrogen; Na<sup>+</sup>: sodium; K<sup>+</sup>: potassium; Cl<sup>-</sup>: chloride.

### Realgar may damage the normal tissue structure of liver and kidney

Compared with 0 mg· kg^−1^ control group, no significant changes in ROW of each organ were observed after administration of realgar for 90 days (Table 5). However, after 60 days of discontinuation, in male rats of the realgar group, the liver ROW was lower than the control rats, and the kidney ROW was higher than that of the control rats (Table 6). All animals were observed during dissection. The organs were normal in texture. No obvious swelling, bleeding, adhesions or ulcers were observed in all organs, thorax, abdomen and pelvis.

**Table 5.** Relative organ weights of male and female rats orally administered realgar for 90 days (n=5/sex/group, 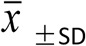)

| Item | Dose (mg·kg<sup>-1</sup>·d<sup>-1</sup>) |  |  |  |
| --- | --- | --- | --- | --- |
|  | 0 | 10 | 40 | 170 |
| **Male** |  |  |  |  |
| **Heart** | 0.27±0.02 | 0.27±0.01 | 0.29±0.01 | 0.26±0.02 |
| **Liver** | 2.32±0.16 | 2.66±0.31 | 2.52±0.29 | 2.53±0.15 |
| **Spleen** | 0.18±0.02 | 0.21±0.03 | 0.21±0.04 | 0.20±0.01 |
| **Lung** | 0.34±0.02 | 0.34±0.02 | 0.35±0.04 | 0.37±0.03 |
| **Kidney** | 0.58±0.04 | 0.61±0.08 | 0.57±0.14 | 0.60±0.05 |
| **Brain** | 0.43±0.03 | 0.41±0.02 | 0.40±0.05 | 0.32±0.18 |
| **Stomach** | 0.04±0.05 | 0.02±0.01 | 0.02±0.00 | 0.02±0.00 |
| **Adrenal gland** | 0.34±0.06 | 0.31±0.03 | 0.36±0.07 | 0.38±0.08 |
| **Thymus** | 0.77±0.06 | 0.71±0.08 | 0.69±0.12 | 0.62±0.23 |
| **Testis** | 0.17±0.04 | 0.16±0.02 | 0.20±0.06 | 0.17±0.03 |
| **Epididymis** | 0.23±0.08 | 0.17±0.02 | 0.15±0.01 | 0.19±0.08 |
| **Prostate gland** | 0.27±0.02 | 0.27±0.01 | 0.29±0.01 | 0.26±0.02 |
| **Female** |  |  |  |  |
| **Heart** | 0.31±0.03 | 0.31±0.02 | 0.38±0.11 | 0.32±0.03 |
| **Liver** | 2.54±0.16 | 2.75±0.21 | 2.63±0.23 | 2.48±0.22 |
| **Spleen** | 0.25±0.03 | 0.26±0.06 | 0.25±0.01 | 0.27±0.03 |
| **Lung** | 0.43±0.01 | 0.42±0.04 | 0.47±0.05 | 0.47±0.05 |
| **Kidney** | 0.64±0.07 | 0.66±0.06 | 0.64±0.07 | 0.63±0.05 |
| **Brain** | 0.64±0.02 | 0.62±0.03 | 0.67±0.08 | 0.62±0.05 |
| **Stomach** | 0.04±0.00 | 0.04±0.00 | 0.04±0.00 | 0.03±0.00 |
| **Adrenal gland** | 0.35±0.14 | 0.29±0.11 | 0.26±0.06 | 0.25±0.03 |
| **Thymus** | 0.07±0.03 | 0.07±0.01 | 0.08±0.01 | 0.06±0.02<sup>a</sup> |
| **Ovary** | 0.21±0.05 | 0.20±0.10 | 0.18±0.04 | 0.17±0.04 |
| **Uterus** | 0.31±0.03 | 0.31±0.02 | 0.38±0.11 | 0.32±0.03 |

Results are shown as the mean ± SEM for 5 rats per sex in each group. Significantly different from 0 mg·kg<sup>-1</sup>·d<sup>-1</sup> control group by ANOVA: <sup>a</sup> $p < 0.05$ , <sup>b</sup> $p < 0.01$ , <sup>c</sup> $p < 0.001$ .

**Table 6.** Relative organ weights of male and female rats after 60 days’ withdrawal(n=3/sex/group, 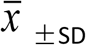)

| Item | Dose (mg·kg<sup>-1</sup>·d<sup>-1</sup>) |  |  |  |
| --- | --- | --- | --- | --- |
|  | 0 | 10 | 40 | 170 |
| **Male** |  |  |  |  |
| **Heart** | 0.26±0.03 | 0.27±0.02 | 0.29±0.02 | 0.28±0.01 |
| **Liver** | 2.56±0.02 | 2.30±0.07<sup>a</sup> | 2.43±0.11 | 2.37±0.06<sup>a</sup> |
| **Spleen** | 0.20±0.02 | 0.18±0.02 | 0.20±0.00 | 0.20±0.02 |
| **Lung** | 0.34±0.07 | 0.36±0.01 | 0.39±0.03 | 0.35±0.01 |
| **Kidney** | 0.58±0.03 | 0.65±0.05<sup>a</sup> | 0.70±0.05<sup>b</sup> | 0.59±0.04 |
| **Brain** | 0.39±0.09 | 0.39±0.00 | 0.45±0.04 | 0.39±0.01 |
| **Stomach** | 0.01±0.01 | 0.02±0.00 | 0.01±0.00 | 0.02±0.00 |
| **Adrenal gland** | 0.28±0.08 | 0.43±0.02<sup>a</sup> | 0.43±0.16 | 0.33±0.11 |
| **Thymus** | 0.45±0.29 | 0.68±0.04 | 0.76±0.03 | 0.66±0.07 |
| **Testis** | 0.14±0.02 | 0.14±0.03 | 0.18±0.01<sup>a</sup> | 0.17±0.03<sup>a</sup> |
| **Epididymis** | 0.16±0.04 | 0.20±0.00 | 0.19±0.02 | 0.16±0.02 |
| **Prostate gland** | 0.26±0.03 | 0.27±0.02 | 0.29±0.02 | 0.28±0.01 |
| **Female** |  |  |  |  |
| **Heart** | 0.34±0.01 | 0.34±0.02 | 0.32±0.02 | 0.34±0.03 |
| **Liver** | 2.64±0.16 | 2.88±0.33 | 2.64±0.19 | 2.76±0.04 |
| **Spleen** | 0.24±0.01 | 0.28±0.03 | 0.26±0.03 | 0.26±0.03 |
| **Lung** | 0.47±0.02 | 0.49±0.03 | 0.47±0.04 | 0.48±0.02 |
| **Kidney** | 0.69±0.08 | 0.70±0.02 | 0.67±0.06 | 0.69±0.05 |
| **Brain** | 0.67±0.05 | 0.69±0.04 | 0.66±0.01 | 0.64±0.04 |
| **Stomach** | 0.03±0.00 | 0.03±0.00 | 0.03±0.01 | 0.03±0.01 |
| **Adrenal gland** | 0.26±0.13 | 0.43±0.16 | 0.34±0.13 | 0.34±0.13 |
| **Thymus** | 0.06±0.01 | 0.06±0.03 | 0.06±0.01 | 0.07±0.01 |
| **Ovary** | 0.25±0.06 | 0.22±0.01 | 0.17±0.07 | 0.26±0.01 |
| **Uterus** | 0.34±0.01 | 0.34±0.02 | 0.32±0.02 | 0.34±0.03 |

Results are shown as the mean ± SEM for 3 rats per sex in each group. Significantly different from 0 mg·kg<sup>-1</sup>·d<sup>-1</sup> control group by ANOVA: <sup>a</sup> *P* < 0.05, <sup>b</sup> *P* < 0.01, <sup>c</sup> *P* < 0.001.

The results showed that at a dose of 10 mg·kg^−1^, which is equivalent to 6 times the maximum clinical dose, realgar has no obvious toxicity to rats’ livers after continuous administration for 90 days. With the increase of the realgar dosage, the liver damage caused by it is aggravated. After 60 days of administration, the hepatocytes of rats in middle-and high-dose realgar group showed mild swelling, degeneration, necrosis and focal interstitial infiltration of inflammatory cells. 90 days after administration, the pathological changes of the liver in the middle and high doses of realgar were significantly aggravated, and there was a statistically significant difference compared with the control group (*p*<0.05). In particular, a few animals exhibited degeneration of flake hepatocytes, eosinophilic degeneration, steatosis, and even bridging necrosis. After 30 days of discontinuation, the high-dose realgar group still had mild swelling and a small amount of inflammatory cell infiltration, and the lesions were statistically different from the control group (*p* <0.05). After 60 days of treatment discontinuation, there was still a small amount of inflammatory cell infiltration and mild hepatocyte swelling in the liver of high-dose realgar group rats, but there was no statistical difference between the realgar group and liver histological group (Fig 2).

**Fig 2.**
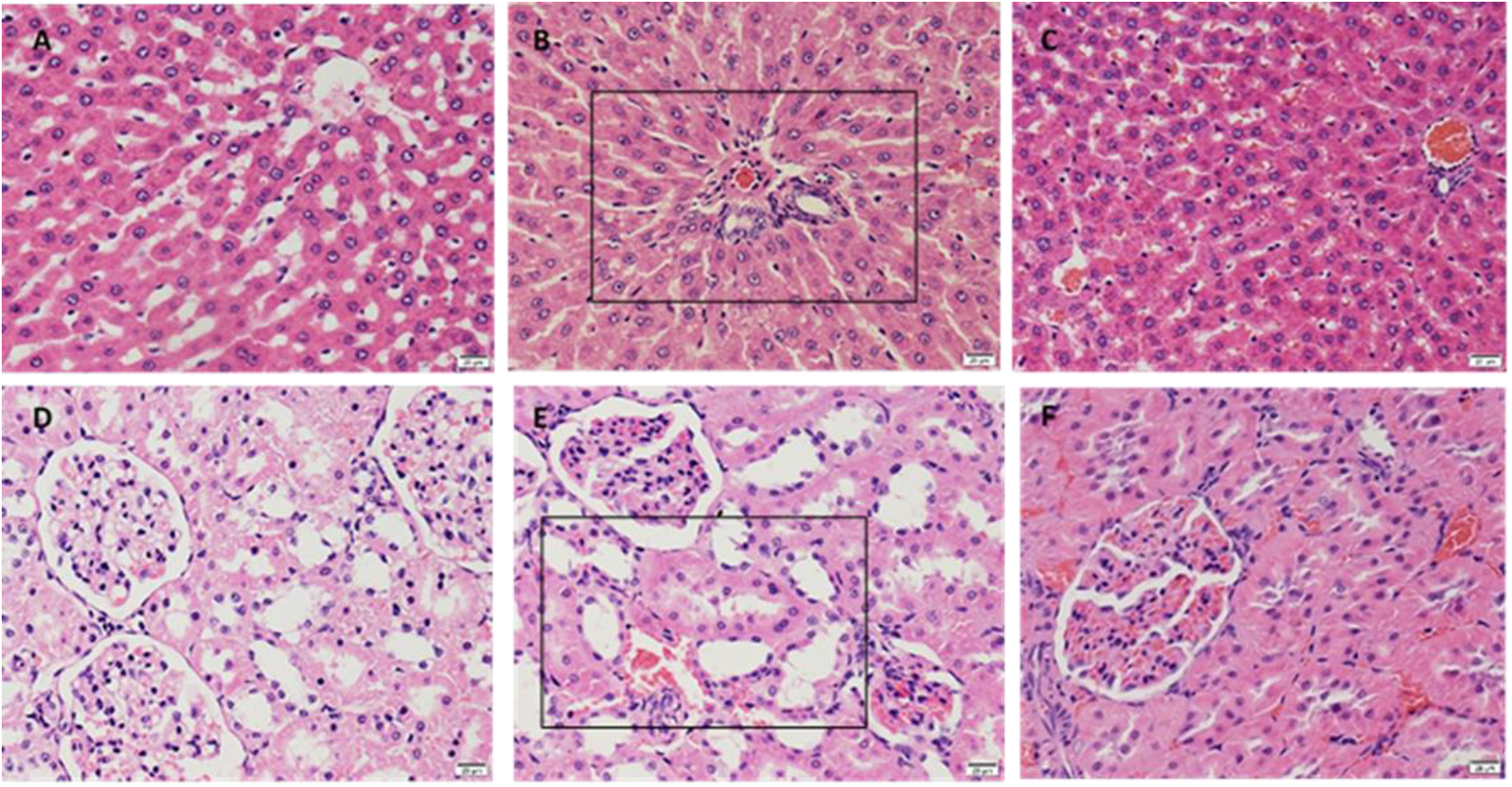
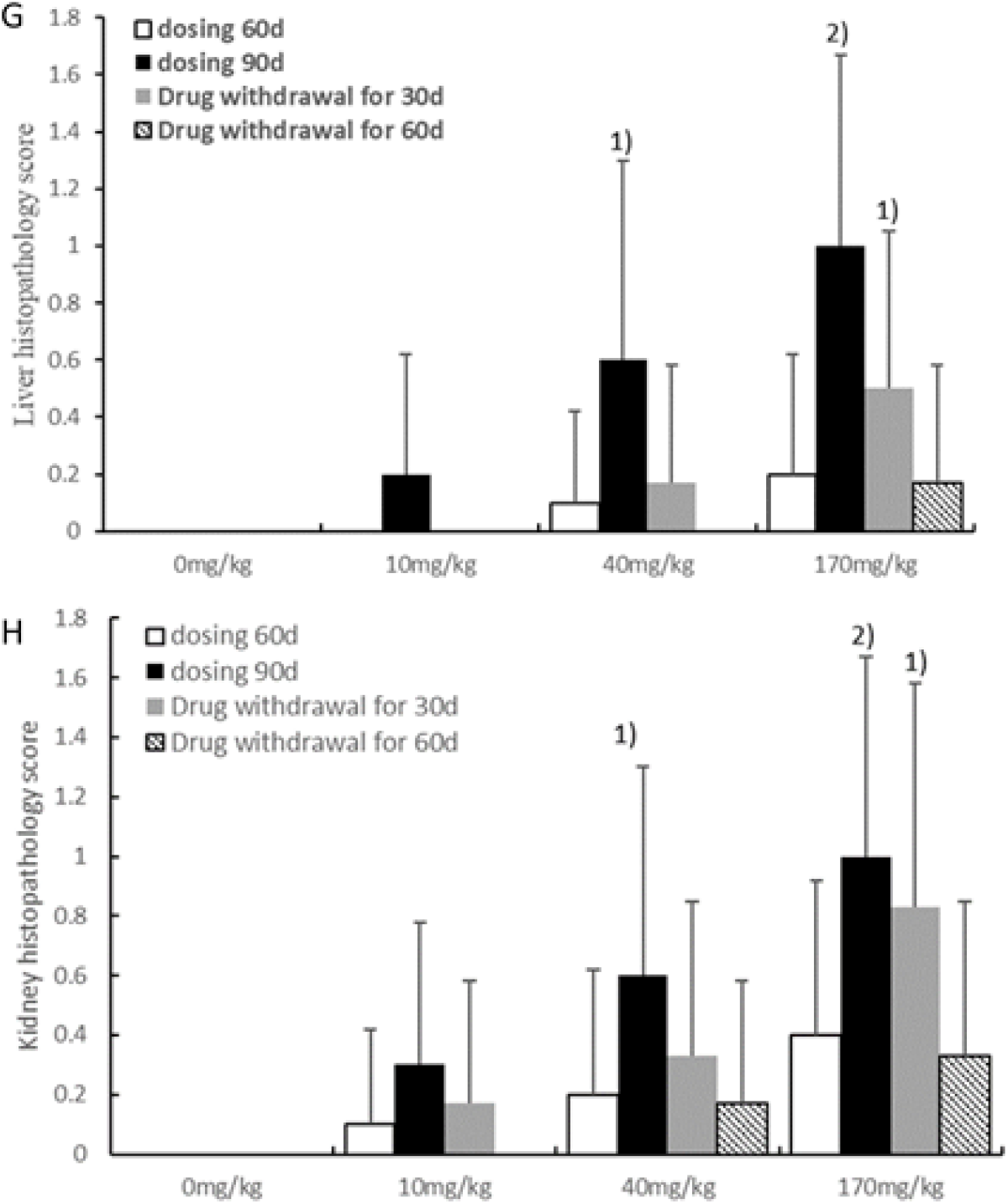
Histopathological examination of liver and kidney sections obtained from rats. (A) After treatment with 0 mg·kg^−1^ of realgar for 90 days, the normal control liver of female rat, (B) After treatment with 170 mg·kg^−1^ of realgar for 90 days, the liver of female rat. The black rectangular box area indicates mild hepatocyte swelling, focal interstitial infiltration of inflammatory cells and intrahepatic bile duct proliferation, (C) 60 days after discontinuation, liver of female ratsthat had been treated with 170 mg / kg realgar, (D) After treatment with 0 mg·kg^−1^ of realgar for 90 days, the normal control kidney of female rats, (E) After treatment with 170 mg·kg^−1^ of realgar for 90 days, the kidney of female rats, (F) 60 days after discontinuation, liver of female rat that had been treated with 170 mg / kg realgar. The black rectangular box area indicates kidney proximal tubule lesion. H&E staining. The bar indicates 20 μm. (G) Liver histopathology damage score and (H) kidney histopathology damage score in four groups. Data are expressed as mean ± SEM. ^1^ *p*< 0.05, ^2^ *p*< 0.01, ^3^ *p*< 0.001. compared with 0 mg·kg^−1^ control group using the Kruskal-Wallis H-test and Mann-Whitney U-test.

The pathological changes of liver tissue were obvious, but there was no obvious abnormality in serum biochemical indicators of reactive liver function. This indicated that the changes of serum biochemical indicators after realgar induced liver injury were later than pathological changes. Thus, when taking realgar for a long period of time, the biochemical indicators commonly used to detect liver function may not reflect early liver injury. Therefore, new sensitive and specific biomarkers reflecting early liver injury should be supplemented. In recent years, the study of drug hepatotoxicity biomarkers has also achieved great development. Several new metabolites, proteins, and nucleic acid microRNAs [26] have been studied as potential biomarkers of liver injury, but still in the experimental research stage. It has been reported that oxidative stress, apoptosis, and up regulation of apoptosis related proteins are the main mechanisms for arsenic induced hepatotoxicity [27-28]. In the large-dose realgar group rats, there was still obvious hepatic injury 30 days after drug withdrawal. Until 60 days after drug withdrawal, the above-mentioned liver tissue pathology basically returned to normal.

After continuous administration of realgar for 60 days in rats, a small number of kidney lesions began to appear. Kidney proximal tubule lesion and mild glomerular lesions were the main lesion, manifested as unclear renal tubular border, luminal narrowing into a star or occlusion and other lesions. After 90 days of continuous administration, the rats in the realgar group showed renal tubular epithelial cells swelling, cytoplasm loosening, vacuolar degeneration, partial proximal epithelial cells necrosis and shedding, the formation of cell tube near the lumen, mild glomerular swelling, congestion and a small amount of inflammatory cell infiltration etc. And as the realgar dose increases, the lesions worsen (Fig 2). These pathological changes were statistically significant compared with the renal tissue of the control group (*p* <0.05, *p*<0.01). After 30 days of withdrawal, the renal lesions in the three doses of realgar group were alleviated, but the renal lesions in the high-dose group were still more severe than those in the control group (*p* <0.05). After 60 days of discontinuation, although one-third of the rats in the high-dose group had slight tubular epithelial swelling and degeneration, there was no significant difference in renal lesions compared with the control group. In conclusion, when the dose of realgar was >10 mg·kg^−1^ and the rats were orally administered for 60 days, pathological damage occurred in the kidney, and pathological damage was exacerbated with increasing dose and/or time of administration. Kidney proximal tubules may be the main target site for realgar kidney toxicity. There is a positive correlation between renal injury and realgar dosage and administration time. Some studies have reported that arsenic can cause oxidative damage to the kidneys [29-31]. The renal damage caused by long-term oral administration of realgar is reversible, with the prolonged withdrawal time, arsenic in the kidney gradually decreased, and kidney damage gradually reduced to recovery.

In addition, the histological and cellular structures of the brain, lungs, bronchus, adrenal gland, pancreas, bladder, testis, ovary, uterus, and other organs of realgar groups had no obvious pathological changes compared with the control group during administration and withdrawal. Some studies have shown that early exposure to arsenic can cause brain cholinergic deficits. Arsenic accumulates in brain astrocytes and affects their viability and glutathione metabolism [32-34]. Due to the limitations of this experimental method, neurotoxicity cannot be directly reflected. Therefore, in this study, there was no direct and concrete evidence that realgar would damage the brain or nervous system of rats (if there are no abnormal behavior changes and abnormal pathological changes, etc.).

### Realgar led to a significant increase in DMA content in tissues

At the end of 60 days’ administration, the total arsenic content (tAs) in rats of realgar groups was significantly higher than that of 0 mg·kg^−1^ realgar control group (*p* <0.01). With the increase of the realgar dosage, the content of tAS in rats also increased significantly. The content of tAs in different biological samples of rats, arranged in descending order of urine> feces> plasm> kidney> liver> brain. The content of tAs in the above tissues of the realgar group was significantly higher than that of the control group. In plasma, arsenic species content from high to low were DMA>MMA>AsIII>AsV. Compared with 60 days after administration, plasma DMA and tAs levels were increased in each of the realgar group after 90 days of administration, but there was no significant difference. After 90 days’ treatment, the accumulation of plasm DMA in the low-, medium- and high dose realgar groups were 11.11, 19.93 and 23.65 times that of the control group, respectively. After stopping the drug, the arsenic content in the rats of realgar group was significantly decreased, and the arsenic content decreased more significantly with the prolonged realgar withdrawal time. However, tAs and DMA content remained significantly higher than the control group even after stopping for 60 days (*p* <0.001). Except for DMA, there was no significant difference in the other arsenic levels in plasma among the three realgar groups compared with the control group (Fig 3).

**Fig 3.**
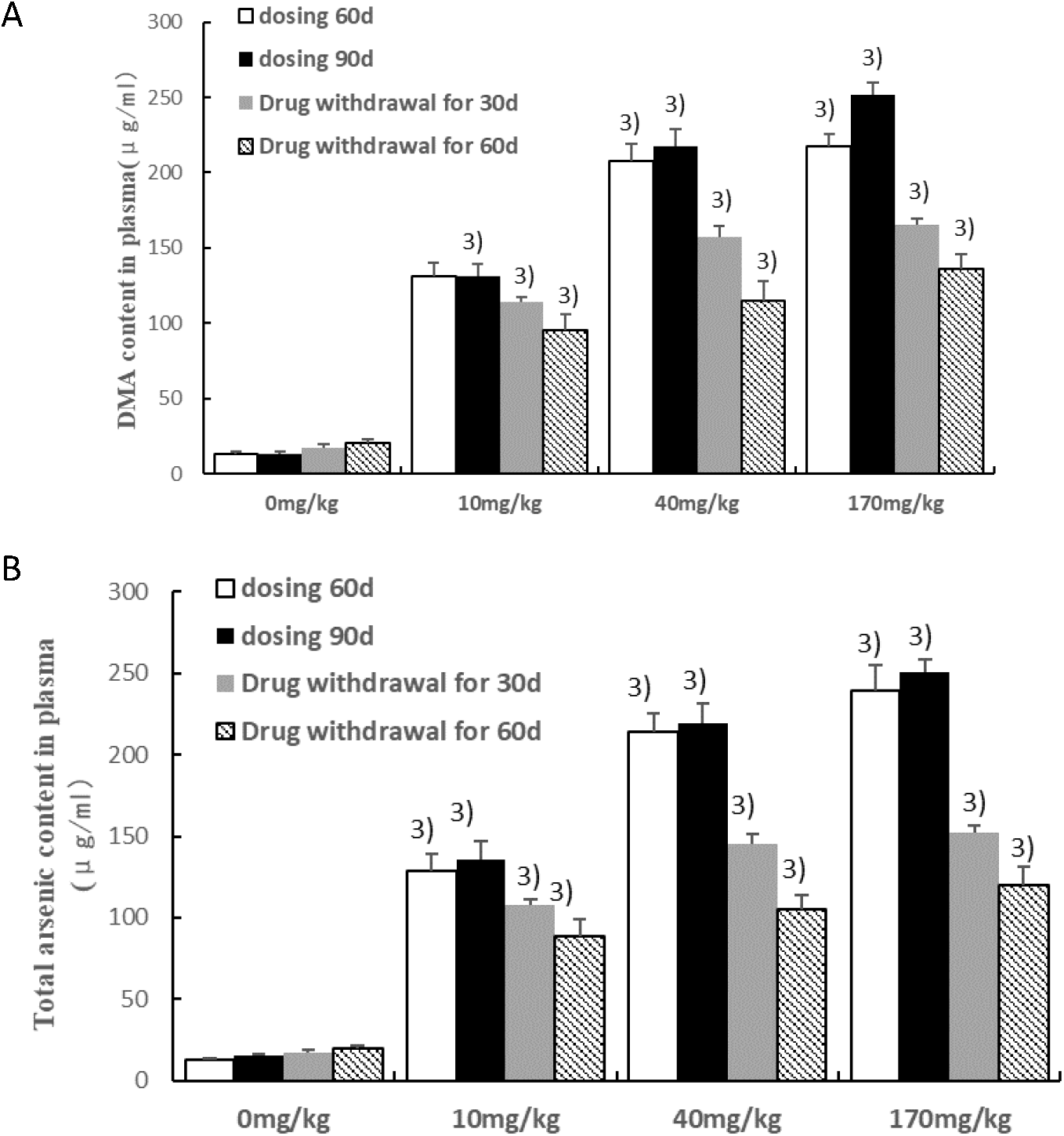
The accumulation of arsenic species in rats’ plasm throughout the experiment. (A) DMA content in rats plasma, (B) Total arsenic content in rats plasm. The content of DMA and tAs in plasma of realgar rats were significantly higher than control rats. Histograms represent average arsenic levels in 10 rats (5 males and 5 females) per group. Data are expressed as mean ± SEM. Significantly different from 0 mg·kg^−1^ control group by ANOVA: ^1^ *p*< 0.05, ^2^ *p*< 0.01, ^3^*p*< 0.001.

The content of tAs in rats’ liver of realgar groups was less than one-tenth the content of the plasma. The accumulation of arsenic in the liver was positively correlated with the realgar dose. The content of arsenic species in the liver of realgar group rats were DMA>AsIII>AsV>MMA>AsB>AsC. After 90 days of oral administration of realgar, the accumulation of DMA in the low, middle and high dose realgar groups were 12.58, 17.75, 28.79 times that of the control group, respectively, which was significantly higher than that of the control group (*p* <0.001), so we conjectured that DMA may play a role in arsenic-induced liver toxicity. There was no significant increase in liver DMA and tAs levels after 90 days of realgar dosing compared to 60 days after dosing. After discontinuation of treatment, all arsenic species levels in the realgar group decreased, while the DMA content was still higher than that of the control group (*p* <0.001) (Fig 4).

**Fig 4.**
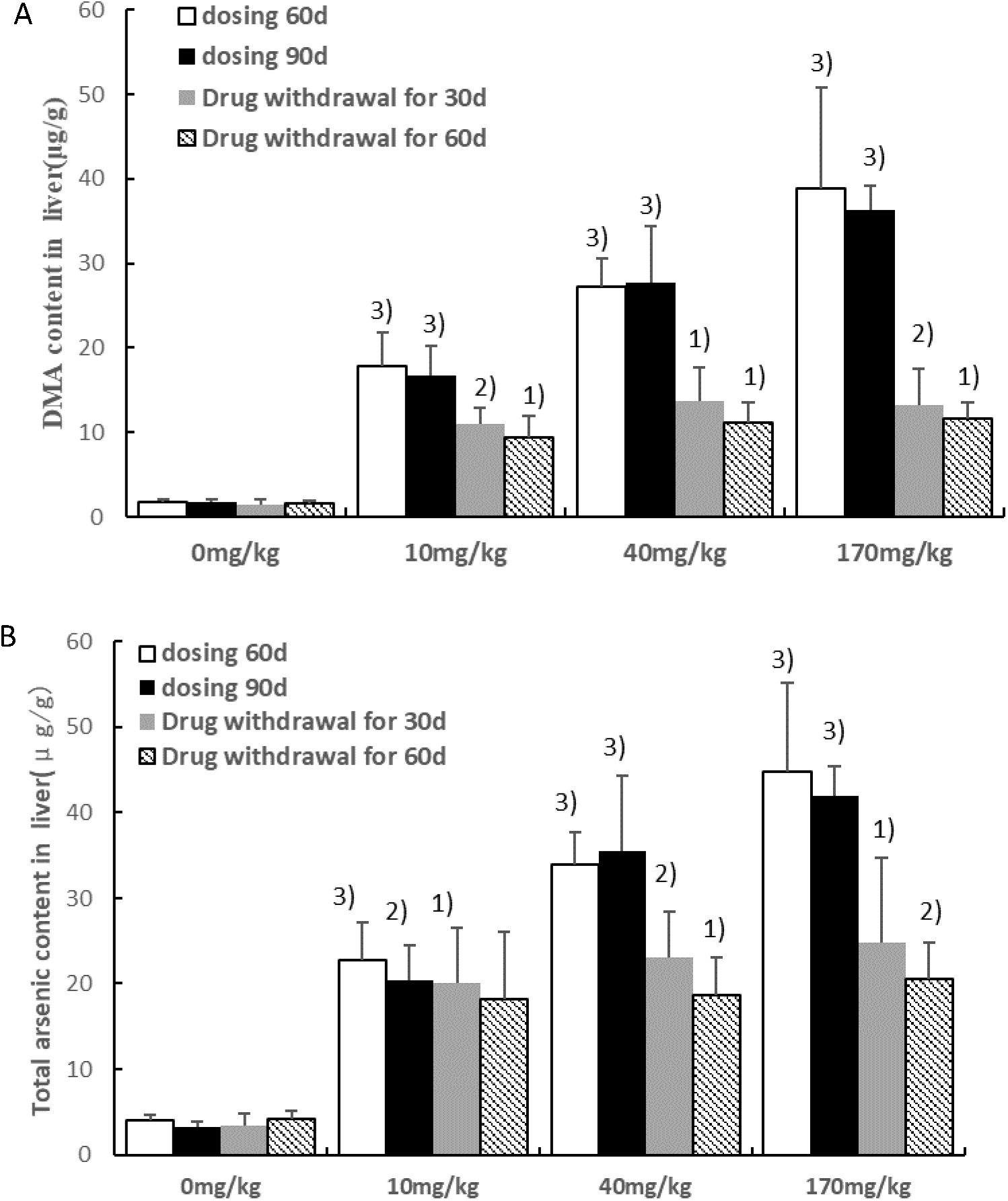
The accumulation of arsenic species in rats’ liver throughout the experiment. (A) DMA content in rats liver, (B) Total arsenic content in rats liver. The content of DMA and tAs in liver of realgar rats were significantly higher than control rats. Histograms represent average arsenic levels in 10 rats (5 males and 5 females) per group. Data are expressed as mean ± SEM. Significantly different from 0 mg·kg^−1^ control group by ANOVA: ^1^ *p*< 0.05, ^2^ *p*< 0.01, ^3^ *p*< 0.001.

The arsenic species content in rat kidneys was DMA> MMA> AsV> AsIII> AsB> AsC. DMA, MMA and tAs levels in the kidney of realgar rats were significantly higher than those of normal rats. (*p* <0.01, *p<*0.05, *p* <0.01), and the levels of DMA, MMA and tAs in the kidneys of rats increased significantly with the prolongation of the administration time. Compared to 60 days after the administration, although the increase of DMA and MMA levels in the kidneys of the realgar-treated rats was not significant at 90 days after the administration, the pathological lesions were significantly aggravated. After discontinuation of treatment, the content of tAs and DMA in the kidneys of the realgar groups gradually decreased, but they were still higher than those of the control group (*p<*0.05). In addition, the renal MMA content also decreased rapidly too, and there was no significant difference compared with the control group (Fig 5). 60 days after drug withdrawal, renal function and renal tissue damage almost completely returned to normal. In view of the above results, we speculate that DMA and MMA in the kidney may be related to arsenic-induced oxidative stress [35] (Zheng, L. Y.et al., 2015), and when they accumulate to a certain concentration, it can induce kidney damage.

**Fig 5.**
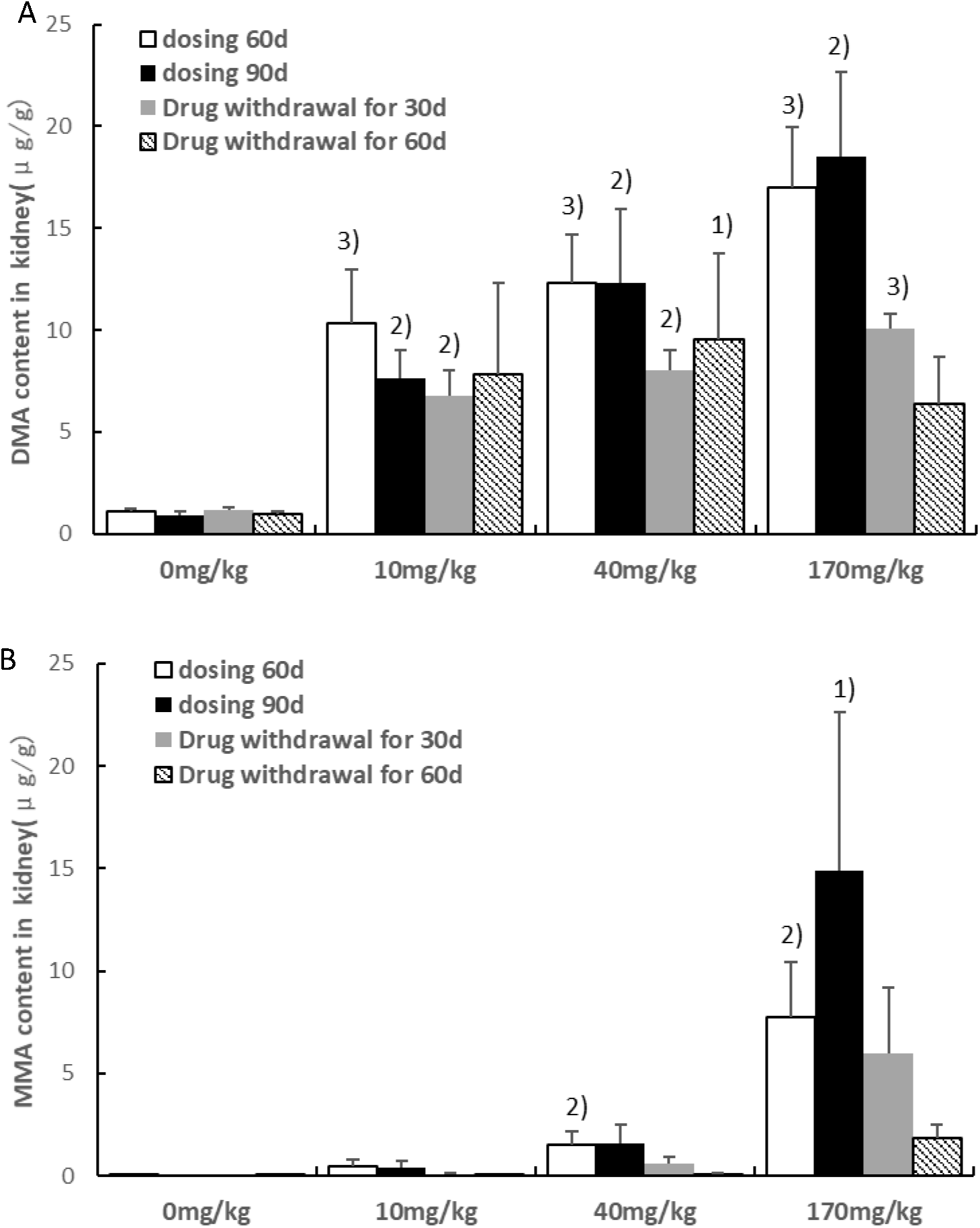
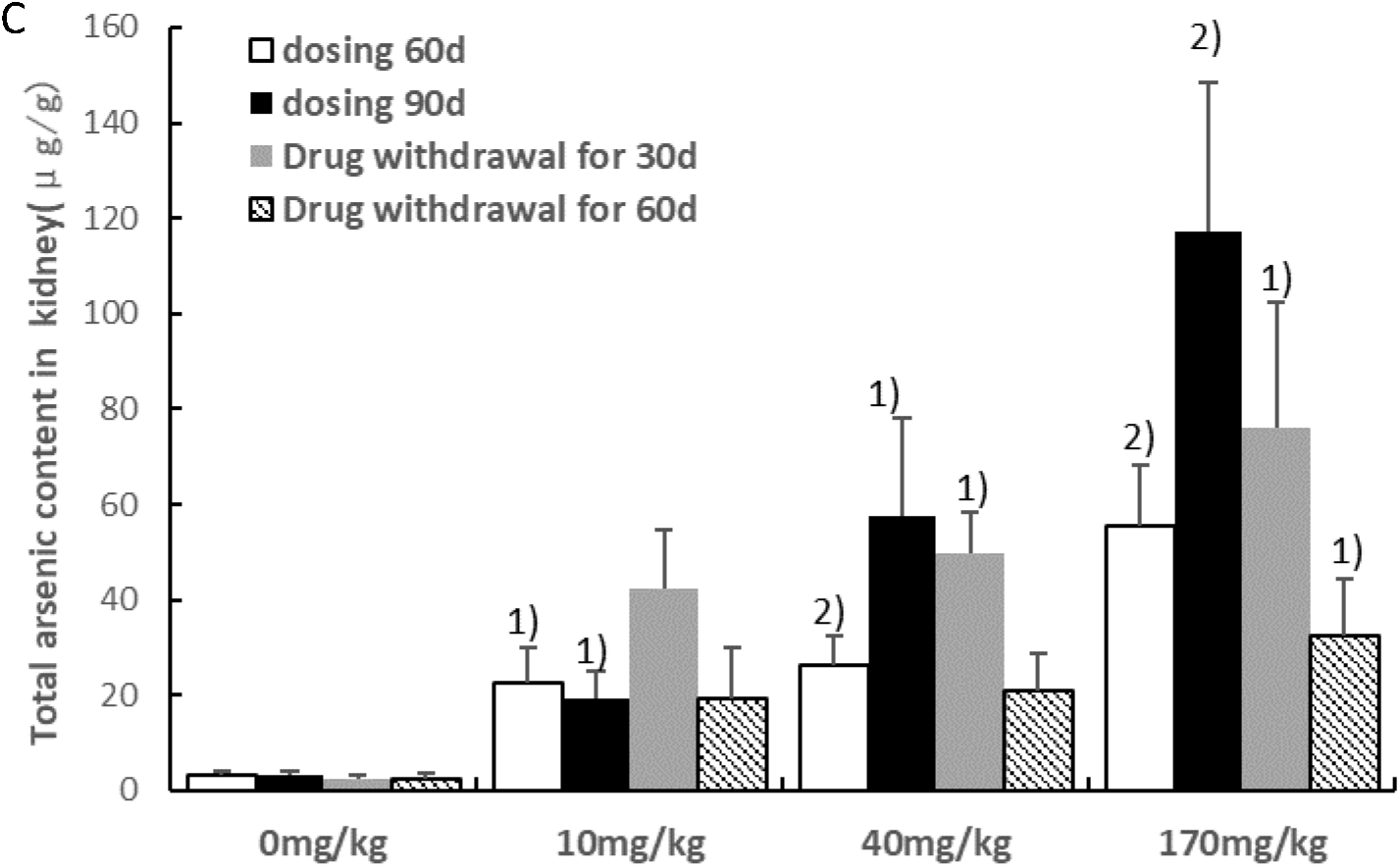
The accumulation of arsenic species in rats’ kidney throughout the experiment. (A) DMA content in rats kidney, (B) MMA content in rats kidney, (C) Total arsenic content in rats kidney. The content of DMA, MMA and tAs in kidney of realgar rats were significantly higher than control rats. Histograms represent average arsenic levels in 10 rats (5 males and 5 females) per group. Data are expressed as mean ± SEM. Significantly different from 0 mg·kg^−1^ control group by ANOVA: ^1^ p< 0.05, ^2^ p< 0.01, ^3^ p< 0.001.

Despite the existence of blood-brain barrier, the content of tAs and DMA in the rats’ brain after realgar administration for 90 days was significantly higher than those in the control group (*p*<0.001, *p*<0.001). This may illustrate DMA can penetrate the blood-brain barrier. Compared with 60 days after realgar administration, the content of tAs in the brain of rats increased 90 days after administration, but the DMA content did not increase significantly. After discontinuation of treatment, tAs and DMA levels in the realgar group began to decrease but remained higher than the control group (*p* <0.05). There was no difference in other arsenic species levels in the brains of realgar and control rats throughout the experiment (Fig 6). Since arsenic is a toxin associated with neurological disease and impaired cognitive function, in order to further understand the relationship between arsenic (DMA) accumulation and brain damage, we believe that a series of in vitro and in vivo tests are needed for research.

**Fig 6.**
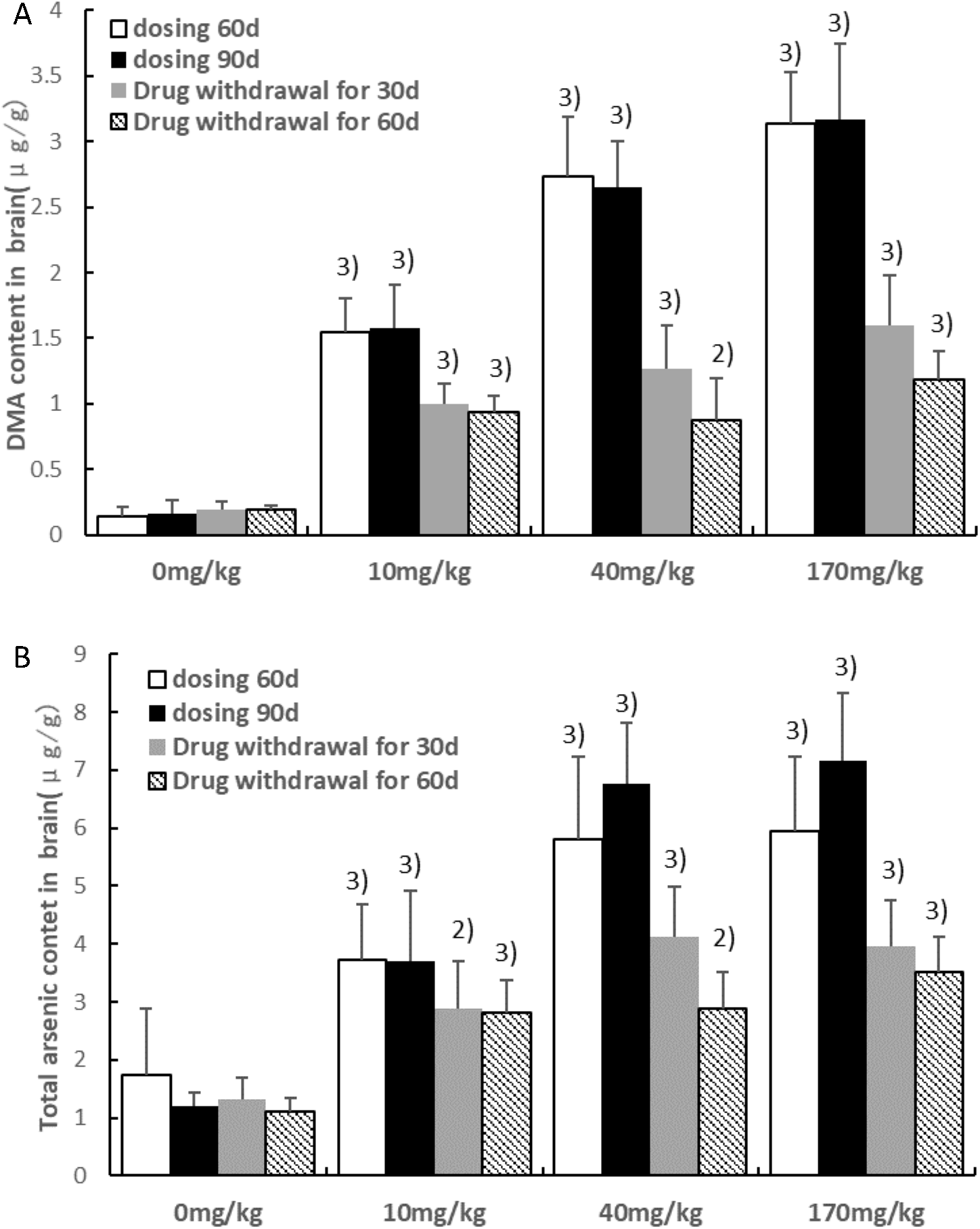
The accumulation of arsenic species in rats’ brain throughout the experiment. (A) DMA content in rats brain, (B) Total arsenic content in rats brain. The content of DMA and tAs in brain of realgar rats were significantly higher than control rats. Histograms represent average arsenic levels in 10 rats (5 males and 5 females) per group. Data are expressed as mean ± SEM. Significantly different from 0 mg·kg-1 control group by ANOVA: 1 p< 0.05, 2 p< 0.01, 3 p<0.001.

The content of tAs in feces and urine of realgar treated group was significantly higher than that of the control group (*p* <0.05). As the dosage of realgar was increased, the tAs in feces and urine of the corresponding rats also increased. In urine, DMA and MMA were the two main arsenic species, and the amount of DMA were obviously higher than MMA. Some reports had shown that high DMA% (and low iAs% and MMA%) in urine represent enhanced methylation capacity which thought to decrease the susceptibility to arsenic-related toxicity [36-37]. Compared with the 60th day after administration, the arsenic content in rat urine increased and the arsenic content in feces decreased 90 days after administration. After realgar stopped for 30 days, the rats in the treatment group were the same as those in the control group, and almost no obvious arsenic was detected in the feces. 60 days after drug withdrawal, the urinary DMA and tAs levels in the realgar group were significantly decreased, but still higher than the control group (*p* <0.05) (Figs 7-8).

**Fig 7.**
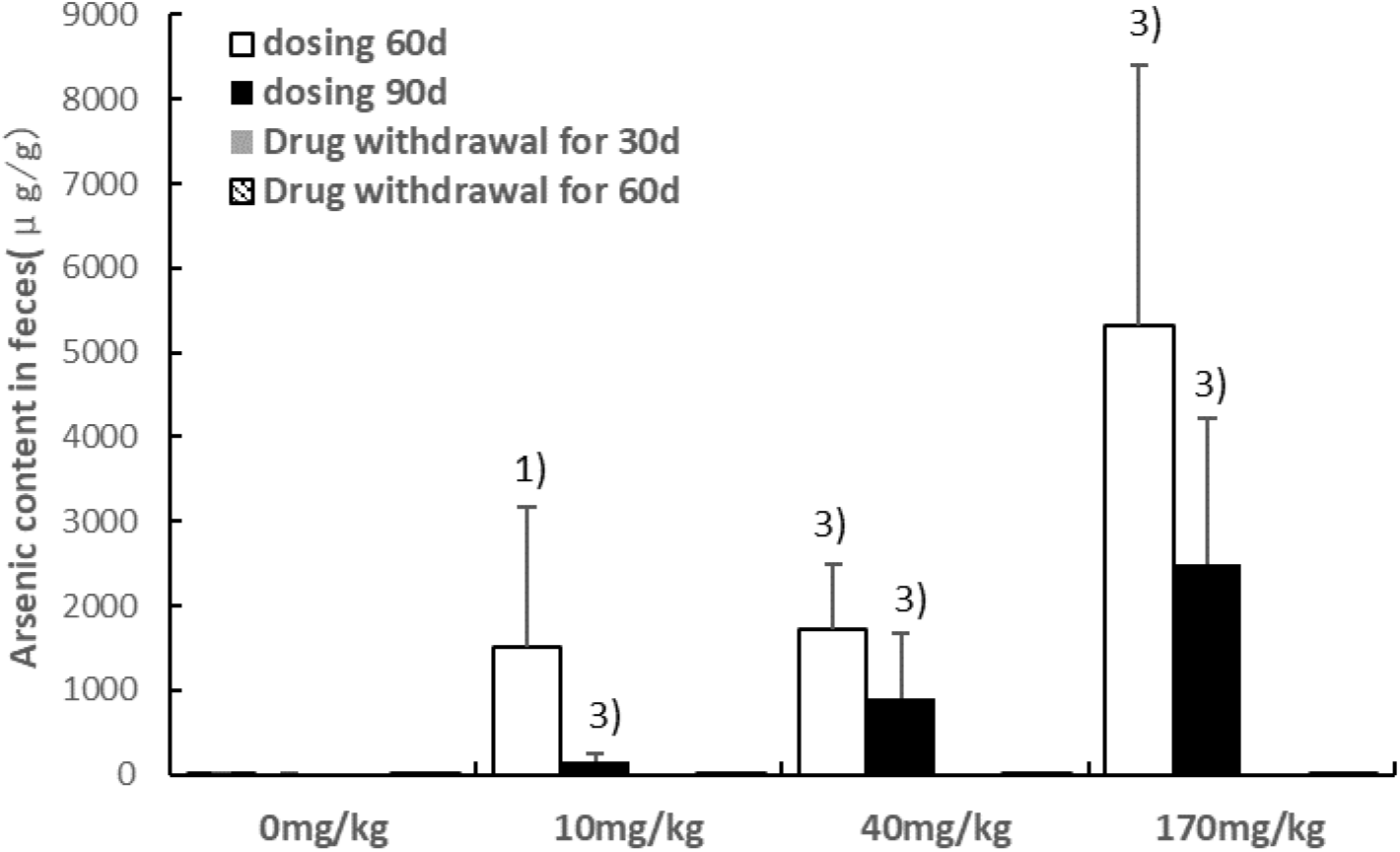
The accumulation of total arsenic in rats’ feces throughout the experiment. The content of tAs in feces of realgar rats were significantly higher than control rats during realgar administration. Histograms represent average arsenic levels in 10 rats (5 males and 5 females) per group. Data are expressed as mean ± SEM. Significantly different from 0 mg·kg^−1^ control group by ANOVA: ^1^ *p*< 0.05, ^2^ *p*< 0.01, ^3^ *p*< 0.001.

**Fig 8.**
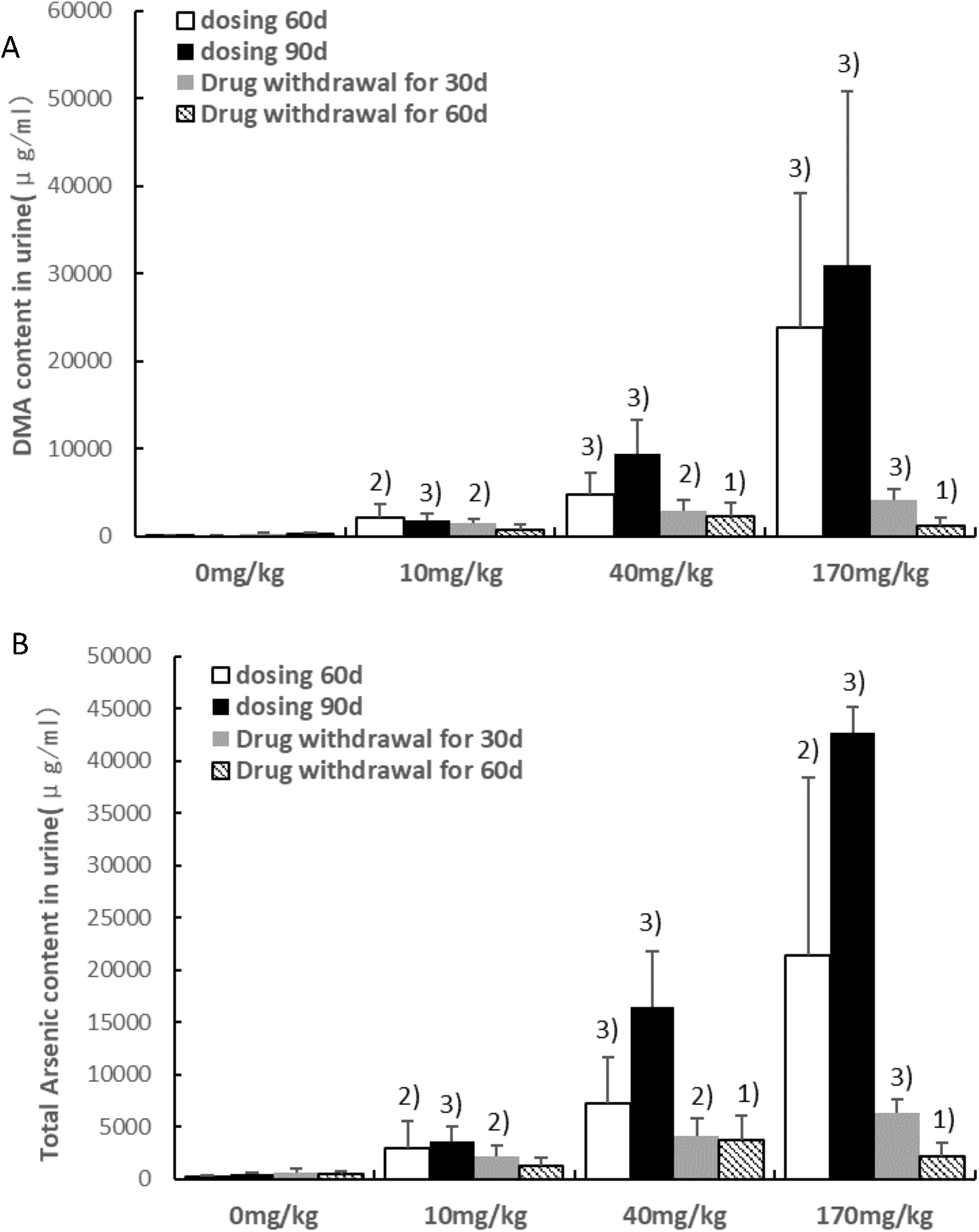
The accumulation of arsenic species in rats’ urine throughout the experiment. The content of DMA and tAs in urine of realgar rats were significantly higher than control rats. Histograms represent average arsenic levels in 10 rats (5 males and 5 females) per group. Data are expressed as mean ± SEM. Significantly different from 0 mg·kg^−1^ control group by ANOVA: ^1^ *p*< 0.05, ^2^ *p*< 0.01, ^3^ *p*< 0.001.

It is known that most of the arsenic species (MMA, DMA, AsIII, AsV, AsB and AsC) are excreted from rat urine and feces after oral administration of realgar. Once discontinuation, the arsenic content in urine and feces of the realgar group dropped drastically, and the decrease was greater than the decrease of arsenic in blood and tissues. After 60 days of discontinued realgar, total arsenic and DMA levels in the blood, liver, brain and kidneys of the realgar rats were still significantly higher than those of the control rats. That is to say, after oral administration of realgar for long time, most of the arsenic species are excreted, but the elimination of arsenic species that accumulated in tissues is relatively slow. This may be one of the reasons that liver and kidney tissues of rats still show pathological damage after drug withdrawal.

## Conclusions

It can be seen from the results of this study that when the dose of realgar is≤10 mg·kg^−1^, continuous oral administration for 90 days does not cause obvious toxicity in rats. While 40mg·kg^−1^·d^−1^ and 170 mg·kg^−1^·d^−1^ of realgar (corresponding to 40-fold and 100-fold the maximum clinical dose, respectively) can cause toxicity in rats, liver and kidney may be the main toxic target organs of realgar. Therefore, the no observed adverse effect level (NOAEL) dose in this study is considered to be 10 mg·kg^−1^. NOAEL is known to be an important reference when setting safety limits for human body equivalent dose (HED). International regulatory agencies recognize a 100-fold safety factor for non-carcinogens. In order to ensure safety, the safety factor for carcinogenic and teratogenic substances is formulated to be more prudent, and there are different regulations depending on different situations. According to the long-term toxicity test guidelines of ICH M3, the clinical safety of drugs with a drug cycle of 2 to 4 weeks can be referred to the 3-month repeated-dose toxicity test of rodents. Therefore, based on the NOVEL dose in this study, the authors preliminarily concluded that the safe use of realgar was approximately 0.1 mg·kg^−1^· d^−1^ for 2-4 weeks of clinical use.

In addition, the kidney and liver are the main organs for arsenic accumulation. The degree of liver and kidney injury and the accumulation of arsenic were positively correlated with the dose and time of administration of realgar. DMA is the main arsenic species after metabolism of realgar, so it may be the main material basis for realgar exerting efficacy and toxicity. Modern studies have shown that DMA exert toxic and anti-cancer effects by inducing cell differentiation and apoptosis. The authors speculated that the content of DMA in tissue has a certain safety range, beyond this range, liver and kidney toxicity will be induced. With the increase of DMA accumulation time or the accumulation of accumulation, the toxic effect will increase. However, the specific quantitative range of DMA-induced toxicity requires more intuitive and accurate tests to determine. Although large doses or long-term use of realgar have the risk of inducing liver and kidney toxicity, this toxic effect is reversible, that is, it can be restored after drug withdrawal. Therefore, it is recommended that liver and kidney function, blood coagulation, fat metabolism and electrolyte levels should be routinely monitored when the realgar preparation is used for more than 2 weeks. Once a toxic reaction occurs, medication should be stopped immediately to restore organ dysfunction.

## Acknowledgments

We thank Ms. Lifang Wang and Mr. Baoqiang Dai (Institute of Chinese Materia Medica, China Academy of Traditional Chinese Medicine, Beijing, China) for assisting with the animal care.

